# Hemin binding causes structural rearrangements in HRI to inhibit activation via autophosphorylation

**DOI:** 10.1101/2024.08.14.607626

**Authors:** Shivani Kanta, Vanesa Vinciauskaite, Graham Neill, Miratul M.K. Muqit, Glenn R. Masson

**Affiliations:** Division of Cellular and Systems Medicine, School of Medicine, University of Dundee, Dundee, DD1 9SY, United Kingdom; MRC Protein Phosphorylation and Ubiquitylation Unit, School of Life Sciences, University of Dundee, Dundee, DD1 5EH, UK

## Abstract

Heme-Regulated Inhibitor (HRI) is one of the four mammalian kinases which phosphorylates eIF2α to facilitate a cellular response to stress through the regulation of mRNA translation. Originally identified for its role as a heme sensor in erythroid progenitor cells, it has since materialised as a potential therapeutic target in both cancer and neurodegeneration. Here we characterise two modes of HRI inhibition of using structural mass spectrometry, biochemical and biophysical techniques. We demonstrate that several ATP-mimetic compounds, including BRAF inhibitors and a compound, GCN2iB, thought to be specific to GCN2, are capable of potently inhibiting HRI. We demonstrate that hemin, a haem-like molecule, inactivates HRI structurally using hydrogen-deuterium exchange mass spectrometry (HDX-MS), and this results in wide-spread structural rearrangement of the protein and how that impacts on the kinase domain through a series of allosteric interactions. This inhibition mainly impacts autophosphorylation, which includes tyrosine phosphorylation, not observed before in the eIF2α kinases.

## Introduction

Heme-Regulated Inhibitor (HRI) was first identified as a key regulator in erythroid cells, where it activates the Integrated Stress Response (ISR) in response to heme deprivation during terminal erythropoiesis^1,2^, facilitating the balanced translation of the alpha-globin and beta-globin chains in response to fluctuating iron levels^2^. The current model of how HRI is activated is thought to be through an inhibitory interaction with heme, which dissociates upon cellular heme depletion, allowing HRI to become active^3–5^. Once activated, HRI phosphorylates its substrate – the alpha subunit of the eukaryotic initiation factor eIF2-eIF2α on Serine 51 (in higher eukaryotes), resulting in the activation of the ISR and the upregulation of the transcription factor ATF4.

More recently, HRI activity has been shown to be a regulator of mitochondrial stress and mitophagy, the autophagic elimination of mitochondria^6–10^. This is thought to occur primarily through the signalling factor DELE1 (DAP-3 binding cell death enhancer 1). Upon mitochondrial stress, the inner mitochondrial membrane protease OMA1 is activated, which in turn promotes the cleavage of DELE1. The resulting C-terminal fragment may then oligomerise and accumulate in the cytosol, which then results in the activation of HRI and activation of the ISR. Furthermore, HRI activity may impact on PINK1 kinase activity, mutation of which is causative for Parkinson’s Disease^11,12^. Currently it is unknown how DELE1 stimulates HRI, and whether this is dependent upon heme release. In additional to mitophagy, it has been shown that HRI also has a role in wider proteostasis. HRI may be activated upon proteasomal inhibition (via compounds such as bortezomib) to facilitate proteasomal-independent protein degradation via the lysosome and the eventual development of resistance^13–15^. Although the exact mechanism of how HRI may be activated within this context is yet to be elucidated.

Altering HRI activity may be beneficial in the context of cancer^14,16–19^, as over-activation of the ISR may lead to highly selective CHOP mediated apoptosis in cancer cells. This has led to the development of compounds capable of activating HRI – namely the 1-((1,4-trans)-4-aryloxycyclohexyl)-3-arylureas (cHAUs) and the N,N’-diarylureas (such as 1-(benzo[d][1,2,3]thiadiazol-6-yl)-3-(3,4-dichlorophenyl)urea, or BTdCPU)-however, their exact mechanism of how they activate HRI has remained elusive, with most studies being conducted within a cellular context ^16,20–23^. Conversely, there may also be therapeutic potential in inhibiting the ISR and HRI in cancer, as constitutive activation of the pathway may promote cancer cell survival ^14,16,17,24,25^.

Given HRI’s role in translation, cancer and neurodegeneration, we have investigated the mechanisms of autophosphorylation, activation and inhibition of human HRI. We were able to demonstrate that HRI is inhibited by the clinically approved RAF inhibitors (RAFi) small molecules Dabrafenib and Encorafenib, but not LY3009120. Additionally, we were also able to demonstrate that GCN2iB, a tool compound used to selectively inhibit the related eIF2α kinase GCN2^26^, is a highly potent inhibitor of HRI. We were unable to demonstrate any direct *in vitro* effect of the small molecule activator BTdCPU – a well characterised activator of HRI in cells, suggesting this compound activates HRI indirectly as had been recently reported^10^. Using Hydrogen Deuterium Exchange Mass Spectrometry (HDX-MS), we characterised the mechanisms of both the inhibitory heme analogue Hemin, and the BRAF inhibitor Dabrafenib, and showed that although they exhibit distinct mechanisms of action, they both ultimately impact on shared aspects of the kinase domain. We show that this inhibition results in a loss of autophosphorylation, including tyrosine residues, which was unexpected. Our data support that heme binding results in a large-scale rearrangement of the protein, with likely additional folding events of disordered segments, and may provide insight into how inhibitors of HRI may be developed in the future.

## Results

### HRI Oligomerization State is Phosphorylation Independent

N-terminally His-Tagged human HRI was purified from *Escherichia coli (E. coli)* cells using affinity chromatography, followed by ion exchange chromatography, and subsequent gel filtration (See Figure 1A) (*4*). Gel filtration (Figure 1B) suggested the protein was homodimeric. SDS-PAGE analysis of the material purified from *E.coli* demonstrated that HRI (monomeric molecular weight: 75017 Da) migrated to >85kDa, suggesting post-translational modification. Treatment with lambda protein phosphatase caused the HRI band to shift back to ∼75 kDa on the SDS-PAGE – this observation being reported previously (*4*, *19*) (Figure 1A).

**Figure 1:**
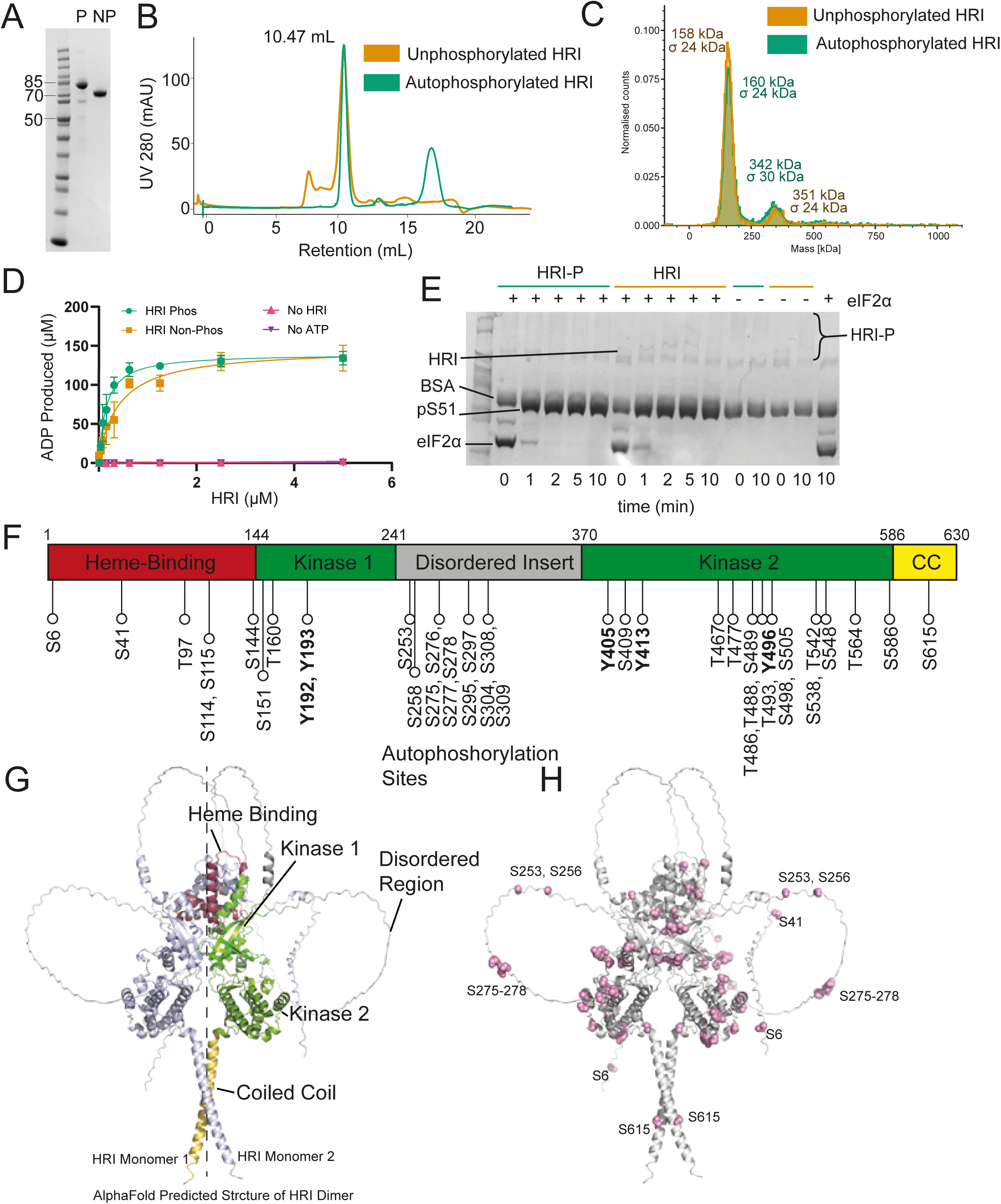
Characterisation of the phosphorylated HRI dimer (A): Purified recombinant HRI run on an SDS-PAGE gel in both phosphorylated (P) and lambda protein phosphate dephosphorylated (NP) protein (B) Size-Exclusion Chromatography of Phosphorylated and dephosphorylated HRI both of which elute with a retention volume of 10.47 mL (C) Mass Photometry analysis of HRI. Both P and NP HRI have a predominant population of ∼150 kDa, indicative of a dimeric oligomerisation state. (D) ADP-GLO Kinase assay of HRI P and NP. (E) Phos-Tag Gel of HRI assay. Both phosphorylation of substrate (recombinant human eIF2α) and autophosphorylation can be seen. (F) Sites of HRI autophosphorylation. (G) AlphaFold 3 model of Apo HRI dimer, with domains highlighted. The dashed line shows the approximate line of 2-fold symmetry. (H) Locations of autophosphorylated residues on HRI, with those on the periphery of highlighted.

We next conducted mass photometry analysis on HRI in both dephosphorylated and autophosphorylated forms to determine whether phosphorylation facilitated any changes in the oligomeric state (Figure 1C). We observed two peaks, a predominant peak with an average mass of approximately 150 kDa, indicative of a dimer, and a second minor peak of a possible quaternary state (with a mass of ∼350 kDa). There was little difference in the relative abundances of these peaks for both dephosphorylated and phosphorylated HRI, suggesting autophosphorylation does not drive dimerization/oligomerisation as previously reported^3^.

### HRI undergoes extensive autophosphorylation

We determined that HRI was extremely active using kinase assays against recombinantly purified human eIF2α (Figures 1D & E). Using Phos-tag gels we were able to track both autophosphorylation and substrate phosphorylation (Figure 1E) – even with nanomolar concentrations of HRI, at very short time points, with the reactions conducted on ice, we found that HRI autophosphorylated rapidly such that the reaction rates against eIF2α between phosphorylated and dephosphorylated HRI were largely indistinguishable (i.e. there was no “lag” autophosphorylation phase). Nonetheless, using the Phos-tag gels, we were able to visualise the gradual autophosphorylation of HRI over 10 minutes.

To investigate the locations HRI autophosphorylation sites, we first dephosphorylated HRI using lambda phopsphatase, removed the phosphatase by gel filtration, and then incubated this dephosphorylated HRI with ATP induce autophosphorylation. Autophosphorylated HRI was then analysed by mass spectrometry to determine sites of phosphorylation. In total 41 sites of autophosphorylation were detected (Figure 1F, Supplementary Data 1, Table 1) – these are sites which could be determined unambiguously in all three repeats and had high stringency criteria. Unexpectedly, we detected numerous phosphotyrosine residues. Crucially we detected the phosphorylation of T486 and T488 – two residues found within the activation loop and previously identified in mice (Mouse residues T483 and T485) as critical for autophosphorylation activity^27^.

**Table 1.**
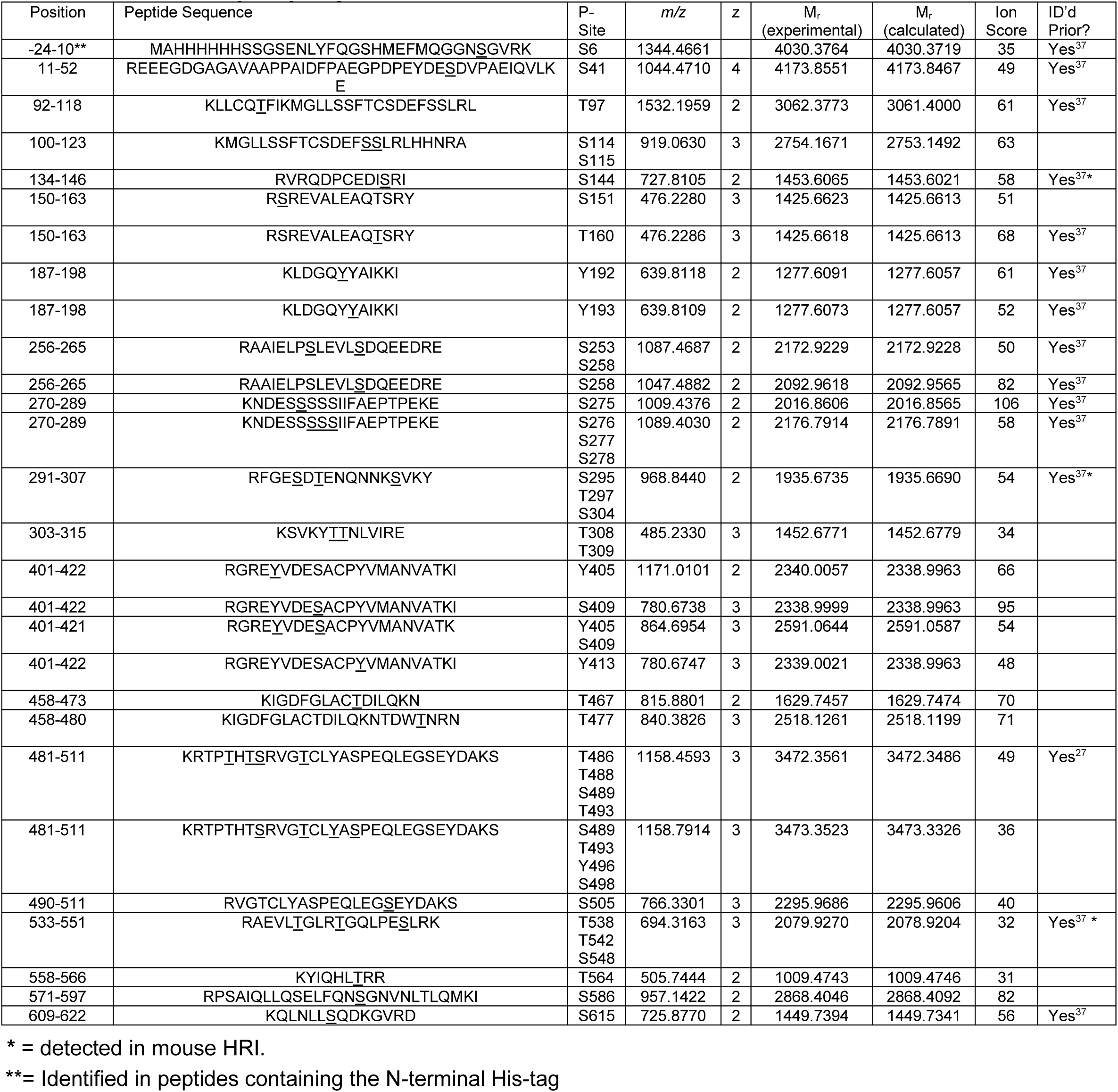
Autophosphorylation sites detected in human HRI.

To interpret the locations of these phosphorylated residues on HRI, we used AlphaFold 3 to create a prediction of the structure of dimeric HRI ^28^ (Figure 1G/H). This prediction placed the kinase domains in a back-to-back conformation, with Heme Binding Domains (HBD) in similar positions “on top” of the kinase domain, with the coiled coil domain situated beneath.

### HRI can be inhibited by RAFi compounds and GCN2iB

Previous studies have used the heme analogue hemin to investigate how iron levels may impact HRI activity^3^. We were able to track both HRI autophosphorylation and eIF2α phosphorylation in the presence of hemin and produced an IC_50_ value of 2.87 µM ± 1.28, which agrees with previous studies^3,29^ (Figure 2A/B/C). We were unable to achieve “complete” inhibition of HRI (as measured by ADP production) using hemin, an observation previously described ^3^, with a residual ∼25% ATPase activity detected even at very high hemin concentrations (>200 μM). Western-blot analysis of HRI found that hemin functionally blocked autophosphorylation, but there was still residual phosphorylation of eIF2α (Figure 2B). We then used differential scanning fluorimetry (DSF) to determine the effects of compound binding on HRI stability (Figure 2F). We were able to detect an HRI unfolding event with a T_m_ of 51.0 ± 0.1 °C and the addition of 5 μM Hemin caused no significant difference to protein stability (50.7 ± 0.3 °C), as has also been previously reported^3^.

**Figure 2:**
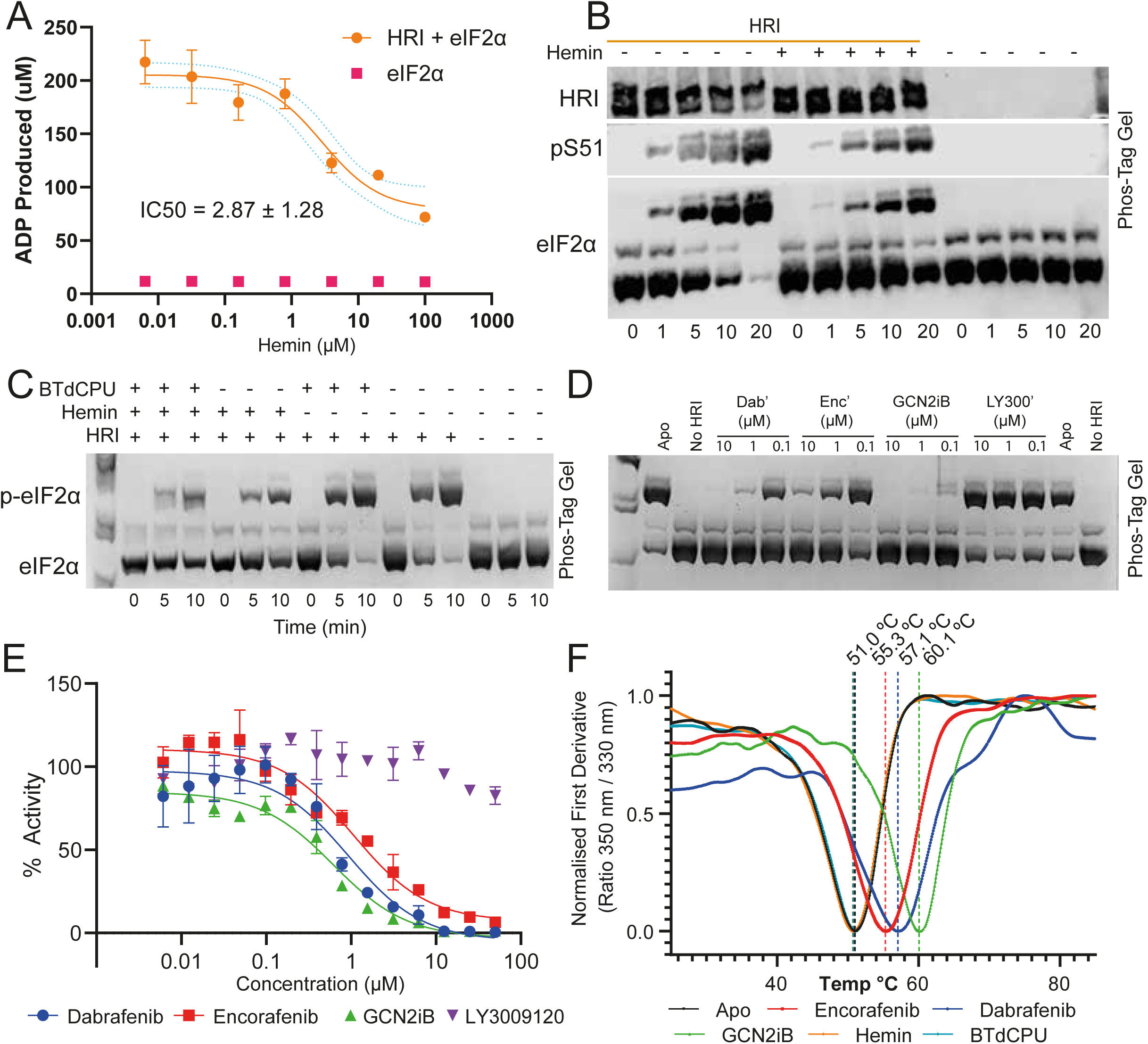
Characterisation of HRI modulators (A) ADP-GLO kinase assay with a concentration gradient of Hemin using recombinant eIF2α as a substrate– an IC50 of 2.87 ± 1.28 µM was determine. No activity was detected in the absence of HRI (eIF2α only control). (B) Western-Blot assay from a Phos-Tag separated HRI kinase assay using eIF2α substrate to observe Hemin effects on both autophosphorylation and substate phosphorylation. The total eIF2α probe shows the development of phosphorylated eIF2α over the course of the reaction, reduced in the presence of Hemin. The S51 phosphorylation specific antibody probe shows the gradual production of phosphorylated substrate from HRI. HRI moves to a higher band (indicative of phosphorylation) upon addition of ATP – this is inhibited upon addition of Hemin. (C) Phos-tag analysis of the effects of Hemin and BTdCPU on eIF2α phosphorylation. (D) Phos-tag analysis of RAFi and GCN2iB. (E) ADP-GLO analysis of HRI activity against RAFi and GCN2iB. (F) nanoDSF analysis of 5 μM Hemin, 10 μM BTdCPU, 10 μM Dabrafenib, 10 μM Encorafenib and 10 μM GCN2iB. TMs (as determined as the nadir of the first derivative of the ratio of the 350/ 330 nm curve) are shown above

It has been reported that small molecule inhibitors of BRAFV600E (RAFi compounds) may have activity against the amino acid stress sensing eIF2α kinase General Control Nonderepessible 2 (GCN2) ^30,31^. Given the sequence homology between the kinase domains of HRI and GCN2 (See Supplementary Figure 1), we screened Dabrafenib, Encorafenib, and LY3009120, and the small molecule activator/inhibitor of GCN2, GCN2iB ^26,32^ to see whether they had any effect on HRI activity (Figure 2D and E). We found that two of the RAFi compounds, Dabrafenib and Encorafenib, were potent inhibitors of HRI, with sub-micromolar IC_50_ values, while LY3009120 had no effect on HRI activity. GCN2iB, was an extremely potent inhibitor of HRI with a sub-nanomolar affinity. Thermal stability analysisusing DSF (Figure 2F) showed significantly increased thermal stability to HRI on addition of 10 μM these compounds – Encorafenib increased the T_m_ by +4.35 ± 0.2 °C, Dabrafenib +6.1 ± 0.8 °C and GCN2iB +9.1 ± 0.4 °C (Figure 2F).

Furthermore, we also screened the small molecule activator of GCN2 BTdCPU – a well-characterised activator of HRI ^17,20–23,33^. Despite a wide range of concentrations and approaches being employed, we were unable to detect any of HRI changes in activity upon BTdCPU incubation (Figure 2C, Supplementary Figure 2). We found that BTdCPU could not overcome Hemin-induced inhibition, and that in the absence of Hemin, could not activate HRI. Addition of 10 μM BTdCPU caused no significant difference on HRI thermostability using nanoDSF analysis (50.8 ± 0.5 °C) (Figure 2F).

### Hemin and Dabrafenib binding result in distinct modes of HRI Inhibition

To gain insight into the mode of action of these inhibitors, we conducted Hydrogen Deuterium Exchange Mass Spectrometry (HDX-MS) in the absence or presence of 100 μM Dabrafenib/Hemin. We produced a peptide map of dephosphorylated human HRI consisting of 218 peptides, with a coverage of 89.3%, and average redundancy of 14.1. HDX-MS of Apo HRI identified that there were several unstructured loops maintained in dimeric HRI between residues 1-70, 143-153, 230-370, 467-495 and 583-593 (Figure 3A) – which corresponded well with the AlphaFold predicted structure (See Table 2.0).

**Figure 3:**
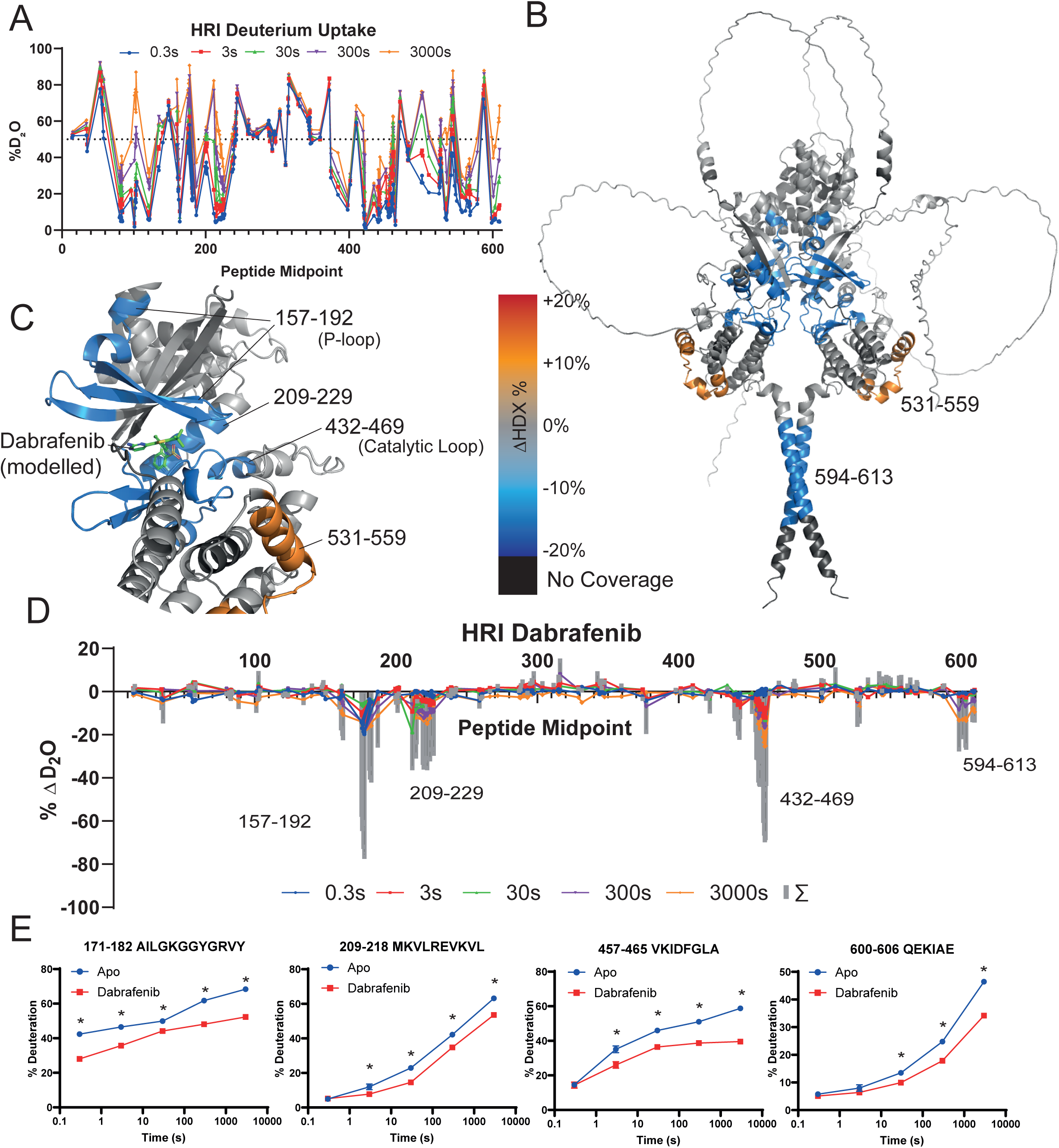
HDX-MS analysis of Dabrafenib Binding (A) Global Exchange profile of Apo Dimeric dephosphorylated HRI over five timepoints. The dashed line at 50% - typically peptides with >50% exchange at the 0.3s timepoint (blue points) are disordered regions (B) HRI kinase domain with Dabrafenib binding modelled. Peptides which exhibited a reduction of solvent exchange upon Dabrafenib binding are highlighted in blue. (C) Inset of the kinase domain of HRI, showing solvent exchange changes upon Dabrafenib binding. Dabrafenib docked using a BRAF model (PDB:5CSW) (D) HDX-MS plot of Dabrafenib binding to HRI. Summed difference over all five timepoints (E) Selected peptides exhibiting differences in solvent uptake upon Dabrafenib binding. * = > 5%/0.5Da difference and passes a student t-test with p = 0.05 (or lower).

**Table 2.** HDX-MS Statistics.

|  | **Apo** | **Dabrafenib** | **Hemin** |
| --- | --- | --- | --- |
| **# of Peptides** | 218 | 218 | 218 |
| **Coverage** | 89.3% | 89.3% | 87.9% |
| **Gaps of Coverage** | 41-46, 66-72, 96-98, 106-113, 366-389, 381-392, 516-521, 615-630 |  |  |
| **Av. Length** | 14.1 | 14.1 | 14.1 |
| **Redundancy** | 4.7 | 4.7 | 5.1 |
| **Time Points** | 0.3 s*, 3 s, 30 s**, 300 s**, 3000 s** |  |  |
| **Replicates** | ND 4, Deuterated 3 |  |  |
| **Controls** | Maximally Deuterated (APO) Control |  |  |
| **Replicability** | 1.08%*** | 0.94% | 0.97%**** |
| **Significance threshold** |  |  |  |

\*= 0.3 s timepoint created by pipetting 3 s by hand at with ice-cold solutions.
\*\*= conducted by liquid handling robot
\*\*\*=Mean standard deviation between replicates
\*\*\*\* = for peptides exhibiting EX2 kinetics only.

Dabrafenib binding (see Figure 3B-E) caused a reduction in solvent exchange in four areas – a kinase inhibitor pocket consisting of residues 157-192, 209-229, 432-469 (Figure 3B, C, D,E) and intriguingly, the C-terminus residues 594-613 (Figure 3B). Single peptides covering the DFG loop – for example, residues 459-464, showed strong reduction in HDX, however neighbouring peptides (*e.g.* 448-456 and 467-475) showed no reduction suggesting a possible direct interaction between Dabrafenib and the DFG loop. Modelling of Dabrafenib in the HRI kinase domain was done using crystal structure PDB:5CSW ^34^, and this aligned well with HDX-MS data. Dabrafenib binding also produced an area of increased solvent exchange in Kinase 2 domain between residues 531-559.

Hemin binding resulted in very widespread alterations in the solvent exchange rate of HRI (Figure 4), suggesting a large structural rearrangement of the protein. Despite the concentration of Hemin being at saturating concentrations (100 μM), many HRI peptides exhibited bimodal distributions upon Hemin binding making data analysis and interpretation challenging (Figure 4B,C, D,E). The N-terminus of HRI which exhibited EX1 kinetics between residues 1-70 on hemin binding (Figure 4B/E)– previously the N-terminus of HRI has previously identified as to undergoing a structural reorganisation upon heme binding ^3,35^. EX1 kinetics were also observed in the disordered region of 144-153, suggesting the possibility of a partial folding event occurring on hemin binding. Furthermore, within the kinase domain, 171-208 also exhibited EX1 kinetics, as well as the very C-terminus of HRI (residues 596-613) addition of hemin – suggesting a widespread conformational change upon hemin binding. The large, disordered region of HRI – between residues 230 and 350, also demonstrated EX1 kinetics, with sub-populations of highly protected peptides emerging on HRI binding.

**Figure 4:**
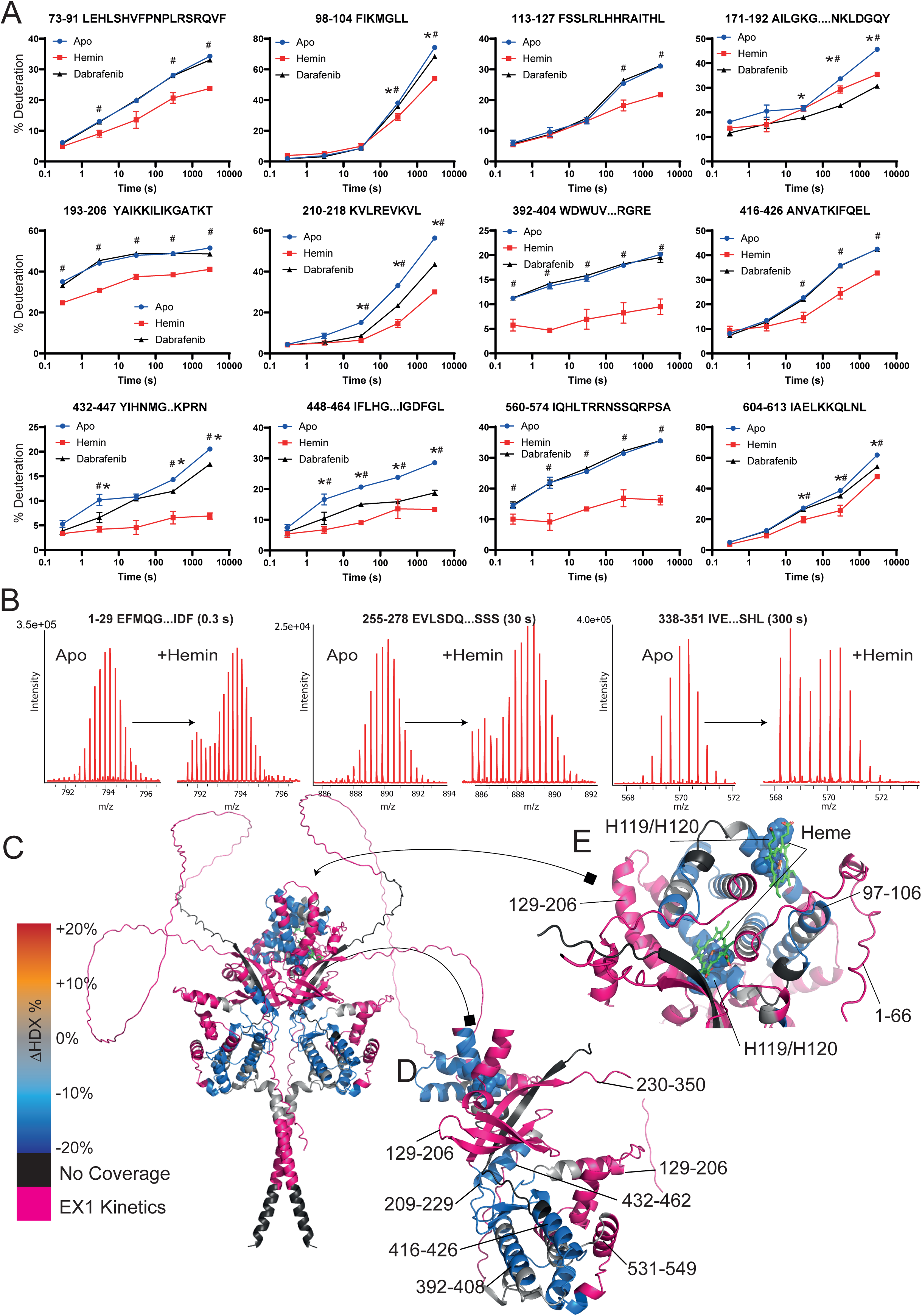
HDX-MS analysis of Hemin Binding (A) A selection of peptides detailing the solvent uptake differences upon Hemin and Dabrafenib binding. * = > 5%/0.5Da difference and passes a student t-test with p = 0.05 (or lower) with Dabrafenib binding when compared to Apo HRI. # = > 5%/0.5Da difference and passes a student t-test with p = 0.05 (or lower) with Hemin binding when compared to Apo HRI. (B) Spectra of selected peptides exhibiting EX1 kinetics which emerge on Hemin binding. (C) HDX-MS data mapped onto an AlphaFold Model with Heme docking. Representations of EX1 kinetics in three peptides found within the regions indicated as pink. (D) Kinase domain upon Hemin binding. (E) HDX-MS data around the Heme Binding Domain. Two previously identified residues, Histidine 119 and 120, are highlighted and represented as spheres.

Several HRI peptides that exhibited “classical” EX2 exchange kinetics on Hemin binding with prototypical isotopic distributions also exhibited decreases– suggesting that incomplete hemin binding was not the root cause of the bimodal peptide populations. Within the kinase domain for example, there is a large reduction in solvent exchange around the activation loop (the “DFG” residues of 461-463) upon hemin binding (Figure 4D), and residues 113-127, which contain the Heme binding histidine residues H119 and H120, exhibited EX2 reductions in solvent exchange upon hemin binding (Figure 4E).

## Discussion

Here we present insight into the activation and inhibition of HRI via biochemical analysis and structural mass spectrometry, providing insights into how heme-like compounds, ATP-mimetic inhibitors, and phosphorylation may work together to regulate this kinase.

The kinase domain of HRI has an unusual architecture, with a very prominent disordered insertion (residues 241-370) between the two lobes of the kinase domain, and a rather extended activating loop (residues 464 to 488), and the P-loop (residues 170-181) lacks a consensus Walker Motif A (typically G-x(4)-GK-[TS])^36^ – HRI has a sequence of _166_EFEELAIL<u>GK</u>GGY_178_ with only the GK resides adhering to the consensus sequence. Opposite the P-loop, the activation loop harbours multiple sites of phosphorylation, including T486 and T488, which have been shown to be crucial for activation of the kinase^27^. Prior to this study, it was unclear how the heme-binding domains of HRI could facilitate the inhibition of HRI. We show in this study that there is likely a large intramolecular rearrangement of HRI caused by Heme binding, that ultimately prevents the kinase domain preventing autophosphorylation.

HRI is an extremely active kinase when released from Heme –experiments had to be conducted rapidly with low nanomolar concentrations of HRI on ice to observe the accumulation of substrate. Furthermore, HRI not only readily phosphorylates eIF2α, but also autophosphorylates rapidly. As many as 33 sites have been reported previously – although these studies were conducted in mice HRI ^37^, here in human HRI we report a total of 41 phosphorylation sites. Although mouse and human HRI have ∼83% sequence homology (see Supplementary Figure 3), some of these sites of phosphorylation are unique to either species. There are two notable patches of dense phosphorylation – a section in the disordered insert (residues 275-309), and within the kinase activation loop of HRI. While phosphorylation of the activation loop is a common means of controlling kinase activation, in HRI we observed the phosphorylation of T486, T488, S489, T493, Y496 and S498, six residues within the 12 amino acids, representing a very dense patch of phosphorylation. Also of note is residue S615 on the coiled coil – this is very distal from the kinase domain and suggests either that this domain is attached to the rest of the enzyme on very flexible loops, or that the placement of the coiled coil is closer to the kinase domain.

Interestingly, we detected numerous phosphotyrosine modifications in the dataset; some have been detected before elsewhere^37^, but the material analysed was purified in a manner which could not exclude the possibility of extraneous kinases modifying HRI. Of particular importance is the residue Y193 (mouse numbering) which was previously found to be critical for the activity of HRI ^37^ – a Y193F mutant had ∼50% reduced activity of wtHRI. Here we identified the phosphorylation of both Y192 and Y193 (although in mice, Y192 is H192, preventing its phosphorylation). Our material, purified from *E. coli,* and subsequently dephosphorylated with lambda phosphatase leaves limited opportunity for these sites to be the result of extraneous phosphorylation. In future work it will be important to assess the stoichiometry of Tyr versus Ser/Thr autophosphorylation using mutagenesis and Phos-Tag analysis.

An example of a protein kinase family that can both tyrosine autophosphorylate and phosphorylate substrates on serine/threonine residues is the DYRK (dual specificity tyrosine-phosphorylation regulated kinase) family. These proteins have been shown to tyrosine autophosphorylate co-translationally and after translation as a mature protein^38^. It would be interesting to perform phosphomapping analysis of HRI in the absence of ATP incubation *in vitro* to determine whether HRI becomes tyrosine autophosphorylated whilst it is inside the *E. coli* cell and further experiments would need to be done to determine whether this can also be a co-translational event.

A key determinant of specificity for serine/threonine versus tyrosine kinases is a single amino acid in a region of the activation segment known as the P+1 loop^39^. In serine/threonine kinases, this residue is a serine or a threonine, and in tyrosine kinases, it is a proline essential for interacting with the phospho-acceptor tyrosine residue. In dual-specificity kinases, for example in the DYRKs, it is a serine^40^, and in HRI T488 is followed by a serine (^481^GKRTPTHT<u>S</u>RV^491^), and several other features of the kinase domain point towards it being a Ser/Thr Kinase (e.g. the APE-6 “GT”, the HRD+2 “K”, and the lack of a doule YY at the DFG+10 position^41^). Other elements point towards Tyr/Kinase, e.g. HRD +4 “RN” ^41^. Notably, the highly conserved “APE” element is found as ApSPE in HRI. Comparing HRI to the DYRK 1A/2/3, many sequences and residues are conserved namely the catalytic loop e.g. a “HRDLKPxNIxL” motif of the catalytic loop and a cysteine at the +2 position of the DFG (DFGxxC) – this cysteine is not present in the other eIF2α kinases (see Supplementary Figure 1). Mutagenesis and further structural studies may help elucidate how the HRI kinase domain is capable of tyrosine phosphorylation.

The discovery that the small molecule BRAF inhibitors Dabrafenib and Encorafenib are capable of inhibiting HRI *in vitro* is of interest. Given that both BRAF and HRI function as back-to-back kinase domains, it may be of interest to determine whether these compounds also exhibit the same concentration-dependent paradoxical activation behaviour as observed in BRAF ^42^. We did not observe activation of HRI by these compounds, but given the extremely high activity of HRI, it may be difficult to detect any additional increase in activity. HDX-MS provides a clear indication of the binding site for Dabrafenib within the kinase domain (Figure 3-B/E), with multiple protections in the kinase site as would expected for a kinase inhibitor. There are reductions in solvent exchange in residues corresponding to both the P-loop (residues 170-183), the catalytic loop (441-449) and DFG motif 461-463) and part of the activating loop (464-508). The two crucial autophosphorylation residues of T486 and T488 showed no reduction in solvent exchange upon Dabrafenib binding – which may mean that these residues may be still capable of being phosphorylated, and potentially “activating” the enzyme while Dabrafenib is still bound.

Additionally, Dabrafenib caused an increase in solvent exchange rate in the kinase domain in residues 531-559. The role of these residues isn’t readily apparent – they may contain a cryptic Walker B motif, although there is considerable variability in this motif. The region does contain resides T538, T542, and S548 all of which are autophosphorylated by HRI. Dabrafenib binding also resulted in the protection in part of the coiled-coil domain in residues 594-613. Given the predicted distance between the kinase domain and this section of the coiled coil, it is difficult to interpret this – other than the possibility that the placement of the coiled coil in the AlphaFold 3 model may be incorrect. This would also explain some aspects of heme binding data (see below)

Hemin binding appears to be an incomplete inhibitor of HRI. This has been reported previously ^3–5^, and here we show that hemin binding appears to be capable of preventing autophosphorylation while still allowing for some eIF2α phosphorylation (Figure 2B/C). This may be due to the fact that hemin is not heme – in a cellular context there will be variety of hemes which have different chemistries with different functional groups, and hemin contains ferric (Fe^3+^) rather than ferrous (Fe^2+^), as found in heme. It may well be that certain hemes are more potent inhibitors of HRI. However, we did find that hemin binding resulted in reductions in solvent exchange at H119/H120 – the proposed haem biding site of HRI ^35^. A further conserved residue C411 (see supplementary figure 3) has been implicated in heme binding also. Although this exact residue did not appear to have a reduction in solvent exchange, the neighbouring helices of 416-426 and 392-408 both had reductions in solvent exchange (Figure 4D). Given the wide-spread EX1 kinetics resulting from large-scale domain reorganisation makes identifying the exact site of heme binding less robust. Overall, previous mutagenesis studies had also suggested that heme binding causes large-scale structural rearrangements of HRI^27,35,37,43^, and this had been accompanied by changes in helicity as measured by circular dichromatism^29^. Our data support this observation – the structure of HRI with heme bound may well be vastly different to that of unbound, autophosphorylated HRI.

Finally, BTdCPU has been widely used as an HRI activating compound ^10,17,33^ – although its mode of action was unclear – previous studies had suggested that N,N′-diarylureas were capable of directly binding and activating HRI^23^. Throughout our studies on HRI, we were unable to detect direct binding or activation with BTdCPU under numerous differing conditions (Supplementary Figure 2). To facilitate HRI activation, the previously suggested model would require BTdCPU to displace a heme molecule to facilitate activation – this would require a K_d_ of < 1 μM, but we were unable to detect any binding or effect on HRI activity even at concentrations greater than 100 μM. This agrees with a recently published study by Perea *et al.* who demonstrated that HRI activation via BTdCPU functioned via mitochondrial decoupling and subsequent OMA1-DELE1 signalling, rather than direct interaction with HRI ^10^.

Given the expanding role of HRI in mitochondrial stress sensing^6–10,12^, it is likely that there will be renewed interest in targeting HRI in disease. Our data here show potential for BRAF-like compounds as inhibitors as a starting point for pharmaceutical development – however there is the distinct possibility that these compounds may also activate HRI via paradoxical inhibitor activation mechanisms ^26,44^. Further structural work is necessary to understand HRI, as current predicted models fail to completely explain the structural changes observed with both heme and inhibitor binding. This study also raises the possibility that HRI may catalyse tyrosine phosphorylation also – mutagenesis and phosphoproteomics work will be necessary to see whether this is maintained *in vivo*.

---

**Supplementary Figure 1:**
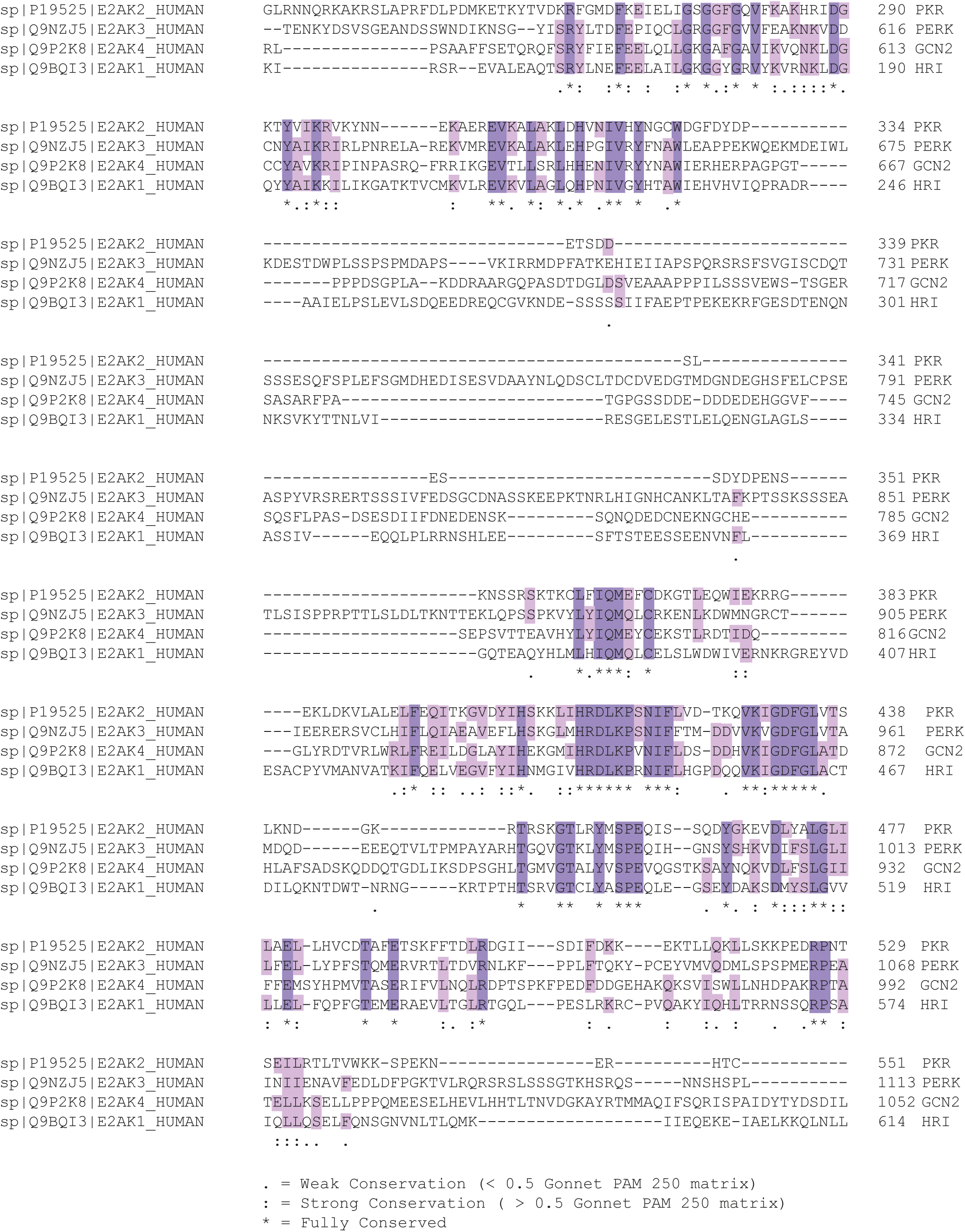
Alignment of the kinase domains of all the human eIF2α kinases (PKR, PERK, HRI and GCN2).

**Supplementary Figure 2:**
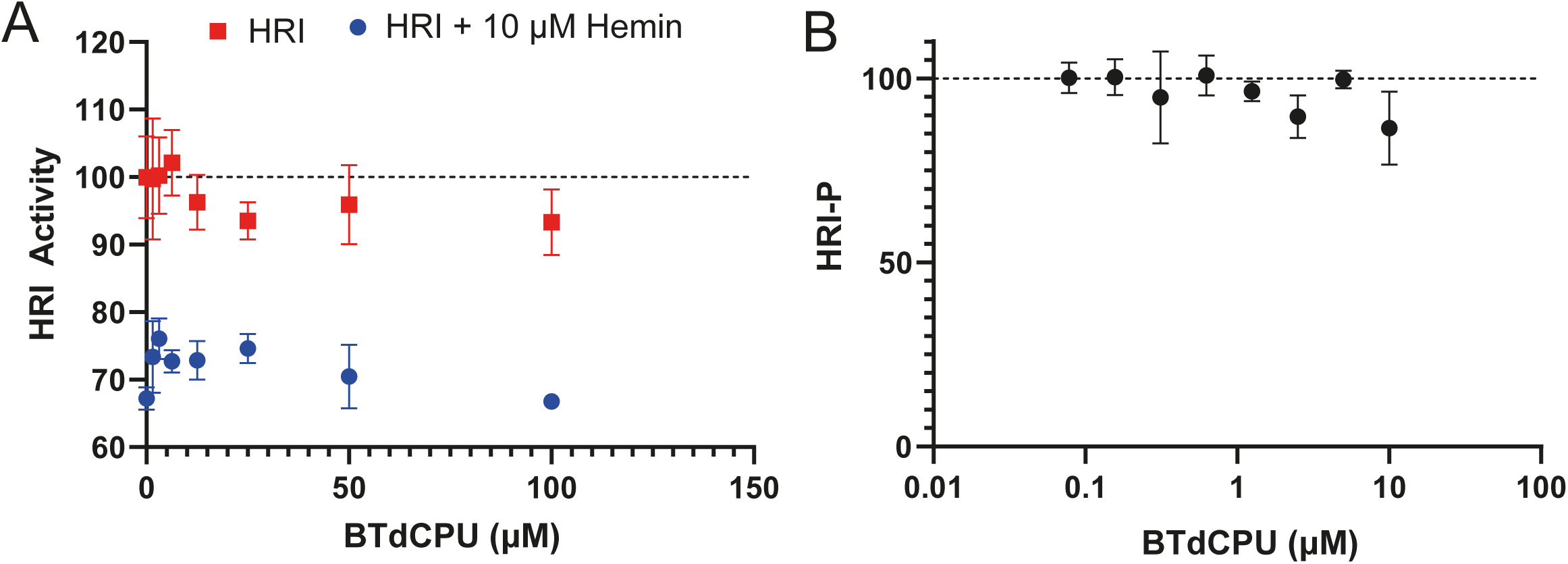
BTdCPU kinase assays. (A) Activity of HRI (as measured by ADP production) on the addition of increasing BTdCPU in either the absence or presence of Hemin. (B) Autophosphorylated HRI (i.e. HRI pre-incubated with ATP and subsequently purified) was exposed to increasing BTdCPU in the presence of eIF2α. ADP levels were measured using ADP-GLO and plotted as a % of APO HRI-P levels.

**Supplementary Figure 3:**
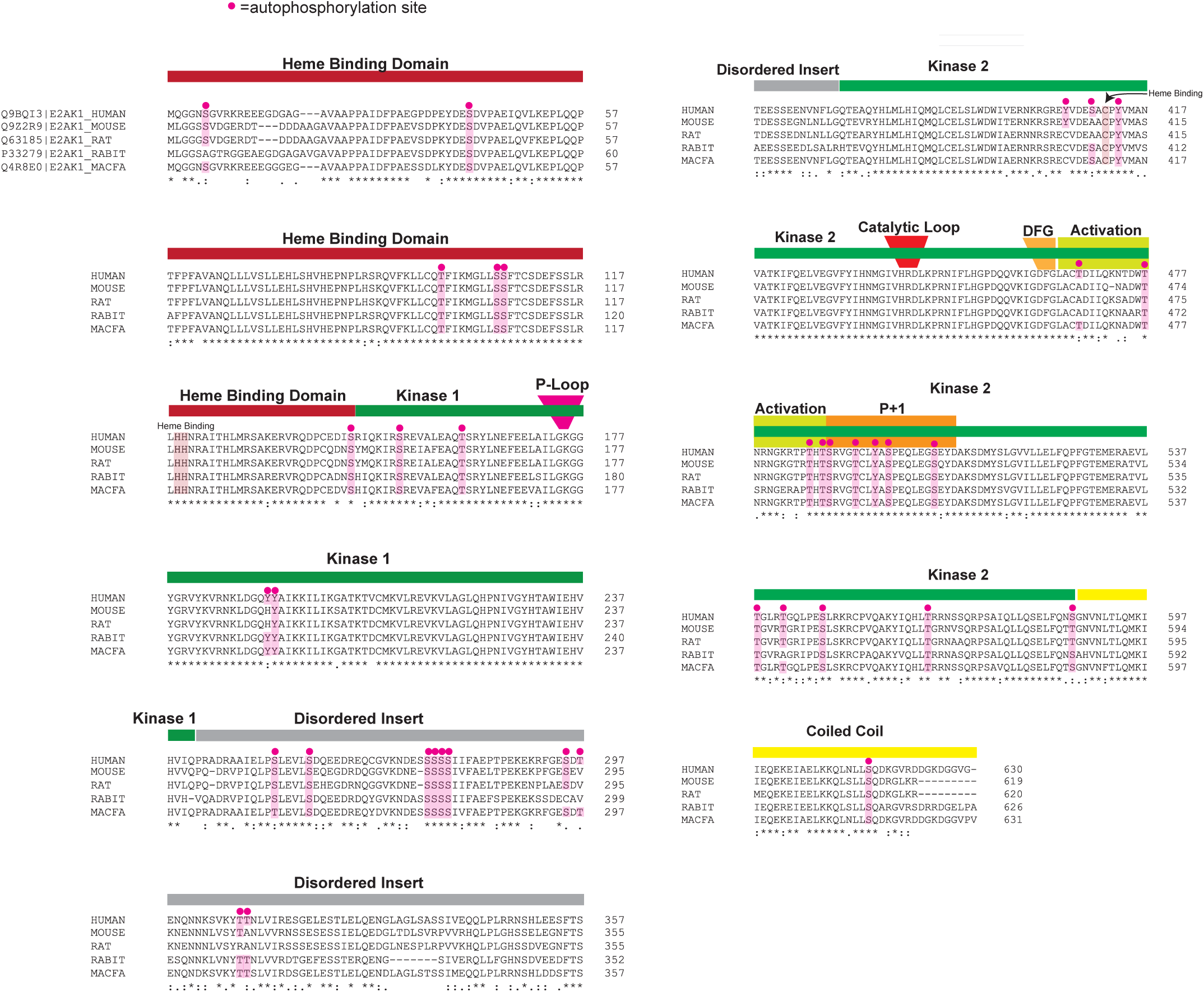
Species alignment of HRI. Human (H. sapiens), Mouse (M. musculus), Rat (R. ratus), Rabbit (RABIT) and Macaca fascicularis (Crab-eating macaque) (Cynomolgus monkey) of EIF2AK1 (HRI).

## Materials and Methods

### HRI Expression and Purification Briefly

Human HRI cDNA was obtained from the MRC Protein Phosphorylation and Ubiquitination Unit (MRC PPU) was subcloned into a pOPTH plasmid with an N-terminal 6His TEV-cleavable tag. Protein was expressed in BL21-Rosetta(DE3) (Novagen) *E. coli*. Cells were grown at 37 °C in 2xTY media until an absorbance of 0.7 at 600 nm was achieved, where protein expression was induced on the addition of 0.8 mM IPTG. Cells were left to express protein for 16 h at 18 °C. A pellet generated from 3 L of media was lysed in 100 mL Lysis Buffer (20 mM Tris pH 8.0, 500 mM NaCl, 20 mM Imidazole, 2 mM beta-mercaptoethanol (BME), with 1 µL benzonase, (Novagen) and 2 EDTA-free Protease inhibitor tablet (PROMEGA)) via probe sonication (10s ON/ 10s OFF at 60% amplitude, on ice). Lysate was centrifuged at 4 °C at 40,000 *g* for 45 minutes. The supernatant was filtered through a 0.45 nm filter and then loaded onto a 5 mL HisTrap HP Column (Cytiva) equilibrated in Lysis Buffer. After washing with 10 column volumes with lysis buffer and a mAU of <100 of achieved, a gradient of NiNTA Buffer B Buffer (20 mM Tris pH 8.0, 500 mM NaCl, 200 mM Imidazole pH 8.0, 2 mM BME) was applied. Fractions were analysed using SDS-PAGE to determine HRI-containing fractions. HRI containing fractions were then diluted 1:1 with Q_O_ Buffer (20 mM Tris pH 8.0, 2 mM BME), and then passed over a 5 mL Q HP (Cytiva) column, equilibrated in Q_A_ Buffer (20 mM Tris pH 8.0, 100 mM NaCl, 2 mM BME), at a rate of 2 ml/min. A gradient of increasing Q_B_ buffer (20 mM Tris pH 8.0, 1 M NaCl, 2 mM BME) was then initiated, with HRI eluting at approximately 600 mM NaCl. HRI containing fractions were then concentrated using a VIVASPIN 10k MWCO Concentrator (Sartorius) until a volume of <1 mL was produced. We then dephosphorylated HRI using Lambda Protein Phosphatase (Lambda PP (New England Biolabs)), with 800 units of phosphatase, and the reaction left at 4 °C for 16 hours. Finally, dephosphorylated HRI was injected onto a Superdex 200i GL 10/30 (Cytiva) equilibrated in Gel Filtration Buffer (20 mM HEPES pH 7.5, 150 mM NaCl, 2 mM TCEP). HRI containing fractions, eluting at ∼10.5 mL, were pooled and concentrated with a VIVASPIN 10k MWCO Concentrator (Sartorius) until a concentration of 1.5 mg/mL was achieved. HRI was then frozen in liquid nitrogen at stored at −70 °C.

### eIF2α Expression and Purification

Expression and purification of recombinant human eIF2α was conducted as described previously^45^. DNA encoding full-length human eIF2α (NCBI reference number: NP_004085.1) was inserted into the vector pOPTH with an N-terminal His6 tag followed by a TEV protease site. The plasmid was transformed into chemically competent BL21 Star (DE3) cells, and cells were grown overnight before being inoculated to a 50 mL starter culture in 2xTY media containing 0.1 mg/mL Ampicillin. The starter culture was incubated at 37 °C for 90 minutes, then 10 mL starter culture was added to 4 x 900 mL 2xTY media containing Ampicillin. Cultures were incubated at 37 °C until the optical density reached 0.7, and then protein expression was induced by the addition of 0.3 mM isopropyl β-D-1-thiogalactopyranoside (IPTG). Cells were grown for a further 3 hours at 37 °C before being harvested, washed with ice-cold phosphate-buffered saline and frozen in liquid nitrogen.

Bacterial cell pellets were lysed in 100 mL Lysis Buffer (20 mM Tris-HCl pH 8.0, 100 mM NaCl, 5 % v/v glycerol, 2 mM BME 0.5 mg/mL Lysozyme (Sigma L6876), 2 U/mL Benzonase, one cOmplete EDTA-free protease inhibitor tablet (Roche 04693132001) per 50 mL of Buffer). Cells were lysed using a probe sonicator for 5 minutes (10 s on/ 10 s off) and then centrifuged at 140,000 *g* for 45 min at 4 °C. The supernatant was filtered through a 0.2 µm syringe filter before being loaded onto a 5 mL HisTrap HP Column (Cytiva 17524801) equilibrated in Ni A Buffer (20 mM Tris pH 8.0, 100 mM NaCl, 5 % v/v glycerol, 10 mM imidazole pH 8.0, 2 mM BME), followed by the elution of protein via a gradient of Ni B Buffer (20 mM Tris pH 8.0, 100 mM NaCl, 5% v/v glycerol, 200 mM Imidazole pH 8.0, 2 mM BME). Protein purification then proceeded as described for HRI. Proteins were concentrated to ∼10 mg/mL and then snap frozen in liquid nitrogen.

### Phos-tag Assays

250 nM HRI incubated with 8 µM eIF2α and 100 µM ATP in Kinase Reaction Buffer (20 mM HEPES pH 7.5, 150 mM NaCl, 5 mM MgCl_2_, 1 mM TCEP) for stated time at either room temperature or on ice. Hemin (Selleckchem S5645) was dissolved in 10 mM NaOH(*4*). Reactions were terminated through the addition of an equal volume of 2 x Phos-tag Loading Buffer (0.1 M Tris pH 6,5, 0.2 M DTT, 4% w/v SDS, 15% Glycerol, 1 mM ZnCl_2_). Precast 17-well Phos-tag gels (SuperSep PhosTag (50 µmol/L) FujiFilm 192-18001/199-18011) were loaded with 10 µL of the reaction/Loading buffer solution in the suggested running buffer (0.1 M Tris Base, 0.1 M MOPS, 0.1% SDS-PAGE, 1 mM NaHSO_3_). Gels were run for 1 hour at 150 V and stained using Quick Coomassie Stain (Generon).

### Western Blots

For Phos-tag Assays, gels were wash three times in transfer buffer (25 mM Tris, 190 mM Glycine, 10% Methanol, pH 8.3) with 10 mM EDTA. Proteins were transferred using wet transfer. All blocking buffers were 4% BSA in TBST. Antibodies used: HRI (Invitrogen 7H3L3, Rabbit, 1:500), Total eIF2α (Santa Cruz Biotechnology, SC-133132, 1:500), pSer51 eIF2α (Cell Signalling Technology 9721S, 1:250), IRDye680 Goat anti Mouse (Licor 926-68070), IRDye 800CW Donkey Anti Rabbit (Licor 926-32213). All images taken using Licor Odyssey FC system and images analysed using ImageStudio. Blots representative of a minimum of three repeats.

### ADP-GLO Kinase Assays

Using a PROMEGA ADP-GLO Kinase assay, HRI was incubated with eIF2α and ATP at the stated concentrations in Kinase Reaction Buffer (20 mM HEPES pH 7.5, 150 mM NaCl, 5 mM MgCl_2_, 1 mM TCEP) for the stated time at room temperature, until the reaction was quenched using the ADP-GLO Reagent. 4 µL kinase reactions were conducted in Corning 3824 wells before quenching and development using the ADP-GLO/Kinase Detection Reagent as per the manufacturer’s instructions. An ADP/ATP curve was created using the kit for calculation of relative ADP concentrations after kinase reaction quenches. Luminescence measured using a Pherastar CLARIOstar (BMG LABTECH). Data analysis conducted using Graphpad Prism 10 (Graphpad). A simple linear regression model for the ADP calibration curve was used to determine the relationship between luminescence and ADP concentration. Non-linear regression analysis (typically Agonist vs. response (three parameters) was used in Hemin/BTdCPU experiments.

### AlphaFold Prediction

The human HRI sequence was used as a basis for AlphaFold 3^28^, with 2 copies being set as a parameter for dimerization. Heme was also included for prediction of Heme binding sites.

### Mass Photometry

Mass determination of solution phase HRI and phosphorylated HRI Hemin was conducted using a Refeyn TwoMP instrument. Experimental data were collected using Grace Bio-Labs Culture Well Reusable 50-3mm DIA x 1mm Depth Gaskets and High Precision glass microscope slides, repeatedly cleaned in ultrapure water and isopropanol and dried using a stream of nitrogen gas. Data were obtained through the collection of mass photometry videos. Prior to HRI data collection, mass calibration was conducted using BSA (66 kDa) and aldolase (160 kDa). By fitting Gaussian functions to ratiometric contrast values obtained from the protein standards using the DiscoverMP v2.5 software (Refeyn), a linear mass calibration was obtained. HRI proteins were diluted to the stated concentrations in 20 µL 20 mM HEPES pH 7.5, 150 mM NaCl, 5 mM MgCl_2_ and 2 mM TCEP, placed on the slide, and then a one-minute mass photometry video was collected. Data was analysed using the DiscoveryMP v2.5 (Refeyn) software.

### Thermal Unfolding Assays

Thermal Unfolding Assays were conducted using a NanoTemper Panta using Prometheus Standard Capillaries (NanoTemper) (PR-C002). Briefly, HRI was defrosted, and centrifuged at 20,000 g for 10 minutes, and then diluted to 0.2 mg/mL in 20 mM HEPES pH 7.5, 150 mM NaCl, 1 mM TCEP Buffer was incubated with Hemin/BTdCPU on ice for 45 minutes before analysis. A gradient of 1 °C/min was applied to the sample starting from 25 °C to 85 °C. Fluorescence at 330 nm and 350 nm was monitored using an excitation wavelength of 280 nm. Data analysis and identification of inflection points/TMs was conducted using the Panta Analysis Software (NanoTemper).

### HRI Autophosphorylation Assay and Mass Spectrometry

Lambda phosphatase treated HRI was produced as detailed above. 1 mg/ mL HRI was incubated with 200 µM ATP (Promega) for 1 hour at room temperature, before being quenched with SDS-PAGE Loading buffer and running on a 4-12% NuPAGE BisTris Gel (Invitrogen) in MES SDS buffer for 45 minutes. Bands corresponding to the gel shifted HRI protein (there was no protein at the lower, non-phosphorylated starting migration) were excised from the gel and sent for mass spectrometry analysis. Briefly, gel bands were washed sequentially in for 15 min at RT in ∼ 200 µL of 1:1 MPW/Acetonitrile (ACN), 100 mM Ammonium Bicarbonate (Ambic), 1:1 Ambic/ACN, then a final wash in ACN before desiccation in SpeedVac for 10 min. Bands were then reduced in 10 mM DTT, 20 mM Ambic for 60 at 56 °C, followed by a wash in 50 mM Indole-3-acetic acid, 20 mM Ambic for 30 min at RT. Bands were then washed in the Ambic, Ambin/ACN, and ACN for 15 min before drying again with the speedvac. Gels were then digested using Trypsin (Pierce) at 12.5 µg/mL, 20 mM Ambic, overnight at 30 °C. Peptides were then extracted via incubation with ACN for 15 min. The gel was then centrigued, and peptides recovered from the supernatant. To this solution 5% formic acid was added and incubated at for 45 minutes. Samples were dried down, and then reconstituted with 10 µL 5% Formic Acid/ 10 % ACN, and then diluted with 40 µL of MPW. Solutions were then analysed by a mass spectrometry using a Thermo QExactive Plus Orbitrap. Data was processed using Mascot^46^. Only peptides with ppm scores <5, Ion Scores >30, and unambiguous MS/MS assignment of phosphorylation modification assignment were included in results. Experiments were conducted in triplicate, and only assignments found in all three repeats are included in the results.

### HDX-MS Sample Preparation

5 µM HRI was incubated with or without 100 µM Hemin/Dabrafenib for 30 minutes in Protein Dilution Buffer (20 mM HEPES pH 7.5, 150 mM NaCl, 2 mM TCEP). 5 µL of this sample was diluted with 55 µL of Deuteration Buffer (20 mM HEPES pH 7.5, 150 mM NaCl, 2 mM TCEP, 1% DMSO, 96.5 % D2O) (final D2O concentration = 85.95%) for timepoints 3/ 30/ 300/ 3000 s, before being quenched with 20 µL of ice-cold Quench Solution (6 M Urea, 2% Formic Acid), and being snap frozen in liquid nitrogen and stored at – 70 °C. A further timepoint, 0.3 s, was achieved by incubating both the sample and deuteration buffer on ice and then conducting a 3 s exchange reaction. Each exchange reaction was conducted independently four times. A maximally deuterated control where denatured HRI was incubated in 98% D_2_O for 24 h was also conducted to correct for back-exchange.

### HDX-MS Data Acquisition

Data acquisition was conducted broadly as previously described (^47^). Samples were rapidly thawed at room temperature and then injected into automated HDX-MS fluidics and UPLC manager system (Waters). Samples were loaded into a 50 µL loop and then subsequently digested using a Waters Enzymate BEH Pepsin Column (Part No. 186007233) in a 0.1 % Formic Acid solution with a flow rate of 200 µL/min at 20 °C. Peptic peptides then flowed onto a Waters ACQUITY UPLC BEH C18 VanGuard Pre-Column at 1 °C (Part No. 186003975). After digestion, the flowpath was changed to elute the peptides via a Waters ACQUITY UPLC BEH C18 1.7 µm 1.0 × 100 mm reverse phase column (Part No. 186002346). A gradient from 0-85% 0.1% Formic Acid/ Acetonitrile, conducted at 1 °C with a 40 µL/min flowrate was used to elute deuterated peptides which were then ionised using an ESI source. Mass spectrometry data were collected using a Waters Select Series cIMS instrument, from a 50-2,000 m/z range with the instrument in HDMSe mode. A single pass of the cyclic ion mobility separator (with a cycle time of 47 ms) was conducted. A blank sample of protein dilution buffer with quench was run between samples, and carryover was routinely checked to be <1% intensity of the prior sample.

### HDX-MS Data Analysis

Peptide sequence identification was conducted using Protein Lynx Global Server (Waters). Minimum inclusion criteria were a minimum intensity of 5000 counts, minimum sequence length 5, maximum sequence length 35, a minimum of 3 fragment ions, a minimum of 0.1 products per amino acid, a minimum score of 5.0, a maximum MH+ Error of 10 ppm. Subsequent analysis and determination of deuteration values conducted using HDExaminer (Sierra Analytics/Trajan). Experimental design, data acquisition, analysis, and reporting are in line with the community agreed recommendations^48^. HDX-MS date were represented on the Alphafold 3 generated structure of HRI or HRI kinase domain using PyMOL Molecular Graphics System, Version 3.0 (Schrödinger).

## Acknowledgements

GRM thanks Dr Anna Plechanovova and Prof. Ron Hay for help with Mass Photometry. GRM thanks Dr Olawale Raimi for helpful input, reading of the manuscript and help with aspects of the investigation. This work was funded by The Royal Society, research grant: RGS\R1\231147. HDX-MS equipment was funded by BBSRC Capital Equipment Fund BB/V019635/1. VV is supported by an MRC iCase Studentship (grant number MR/R01579/1). This work was supported by the Michael J. Fox Foundation (M.M.K.M). We thank the MRC PPU Reagents and Services facility (MRC I PPU, College of Life Sciences, University of Dundee, Scotland, mrcppureagents.dundee.ac.uk) for the reagents and/or services indicated in this publication. We thank Hajara Lawal of the Fingerprints proteomics services, School of Life Sciences, Dundee, for their help with phosphorylation analysis of HRI, and Douglas Lamont for his continued support of the HDX-MS facility.

## CRediT Author Contribution

**Shivani Kanta,** Investigation **Vanesa Vinciauskaite**: Resources, Writing — review and editing **Graham Neill:** Investigation **Miratul M.K. Muqit:** Writing — review and editing. **Glenn R Masson**: Conceptualisation, Resources, Supervision, Investigation, Funding acquisition, Methodology, Writing — original draft, Project administration, Writing — review and editing.

